# Hippocampus Modulates Natural Sound Processing at Early Auditory Centers

**DOI:** 10.1101/2022.05.11.491576

**Authors:** Eddie C. Wong, Xunda Wang, Ed X. Wu, Alex T. L. Leong

**Author notes:** Correspondence should be addressed to Ed X. Wu, Ph.D. and Alex T. L. Leong, Ph.D.: Laboratory of Biomedical Imaging and Signal Processing, Department of Electrical and Electronic Engineering, The University of Hong Kong, Pokfulam, Hong Kong, Hong Kong SAR, China. &. **Author Contributions:** E.C.W., A.T.L.L., and E.X.W. designed research; E.C.W. performed research; E.C.W., X.W., A.T.L.L., and E.X.W. analyzed data; X.W. provided technical assistance; and E.C.W., A.T.L.L., and E.X.W. wrote the paper.

## Abstract

Despite its prominence in learning and memory, hippocampal influence in early auditory processing centers remains unknown. Here, we examined how hippocampal activity modulates sound-evoked responses in the auditory midbrain and thalamus using optogenetics and functional MRI (fMRI) in rodents. Ventral hippocampus (vHP) excitatory neuron stimulation at 5 Hz evoked robust hippocampal activity that propagates to the primary auditory cortex. We then tested 5Hz vHP stimulation paired with either natural vocalizations or artificial/noise acoustic stimuli. vHP stimulation enhanced auditory responses to vocalizations (with a negative or positive valence) in the inferior colliculus, medial geniculate body, and auditory cortex, but not to their temporally reversed counterparts (artificial sounds) or broadband noise. Meanwhile, pharmacological vHP inactivation diminished response selectivity to vocalizations. These results directly reveal the large-scale hippocampal participation in natural sound processing at early centers of the ascending auditory pathway. They expand our present understanding of hippocampus in global auditory networks.

## Introduction

In the central auditory system, auditory input from the ear transmits to the inferior colliculus (IC), medial geniculate body (MGB) in thalamus, and auditory cortex (AC) along the ascending auditory pathway (1–3). Information is hierarchically relayed along this ascending pathway, as distinct auditory features like amplitude and frequency are gradually extracted and processed throughout each auditory center (3–7). Existing functional frameworks describing auditory processing in the ascending auditory pathway are often examined using basic stimuli such as pure tones and broadband noise (8–10). However, neural representation of simplified acoustic stimuli may not reliably predict responses to natural sounds (8, 11–13), such as vocalizations, which are critical for facilitating communications and behavioral responses (14–16). Natural sound processing requires decoding complex spectrotemporal dynamic properties (17, 18) and additional input from higher-order regions is needed to facilitate tacitly assumed auditory functions such as communication, learning, and memory processes (19–22). Despite the current consensus on the pivotal roles played by AC corticofugal projections in natural sound processing (7, 23–25), emerging structural evidence has revealed that auditory midbrain and thalamus project to non-auditory regions such as superior colliculus (26) and striatum (27), respectively, and receive afferents from sensory, prefrontal, and limbic regions (28, 29). These findings suggest that information can transmit in and out of early auditory centers in the ascending pathway to cortex and beyond, parallel with those at the AC level in the processing hierarchy. We speculate that the auditory network for natural sound processing is far more brain-wide than presently known.

Given its roles in memory, emotion, and learning functions (30–32), we contend that the hippocampus is a strong candidate to participate in brain-wide auditory processing of natural sounds. Notably, the hippocampus has been indirectly linked with auditory processing (33, 34).

Functional studies indicate interactions between the hippocampus and auditory cortex during learning and memory processes. Electrophysiology studies demonstrate that the hippocampus actively engages the auditory cortex to transform auditory inputs into long-term memories that are subsequently consolidated in cortical networks (35, 36). Meanwhile, studies show that specific hippocampal neurons only respond to sounds associated with a trained sound behavioral task (22, 36), implying that the hippocampus participates in the interpretation of complex auditory inputs. Anatomically, the hippocampus can receive and relay auditory signals via reciprocal projections directly with AC (37, 38) and indirectly through parahippocampal regions (38, 39) and forebrain pathways (19, 40), such as the entorhinal cortex, amygdala and medial septum complex. Tracing studies also indicated indirect projections, albeit scarcer, from the hippocampus to the IC and MGB via parahippocampal regions and amygdala (28, 29, 41). However, existing studies have not directly examined the role of the hippocampus in processing auditory inputs at these early auditory centers. Further, most studies so far focus on the cortex. They provide little to no evidence of the possible functional interactions between the hippocampus and auditory regions at the midbrain and thalamic levels. At present, whether and how the hippocampus functionally influences auditory responses, especially in the early ascending auditory centers, remains unknown.

Here, we posit that the hippocampus participates in natural sound processing at early sound processing centers within the ascending auditory pathway, especially the IC and MGB. The ventral hippocampus (vHP) plays a role in processing sensory inputs with an emotional context (42, 43), thus it may influence natural sound processing throughout the ascending pathway. In this study, we examined whether optogenetically evoked vHP activity modulates sound processing in the auditory midbrain, thalamus and cortex. Using a combined optogenetic cell-specific stimulation of Ca^2+^/calmodulin-dependent protein kinase IIĮ (CaMKIIĮ)-expressing vHP neurons and whole-brain functional MRI (fMRI) visualization, we assessed blood-oxygenation-level dependent (BOLD) fMRI responses to two different categories of sound ± natural sound (i.e., vocalizations) and artificial/basic acoustic stimuli. We revealed that the vHP activity enhances auditory responses to vocalizations, but not artificial stimuli or noise, in the IC, MGB and AC.

## Results

### Brain-wide propagation of neural activity initiated at ventral hippocampus

We characterized the downstream targets of the ventral hippocampus (vHP) using optogenetics to selectively stimulate CaMKHα-expressing vHP excitatory neurons, primarily in the dentate gyrus of vHP. Anatomical MRI scans confirmed the location of the virus injection and optical fiber implantation in vHP of all animals (**Figure 1A**). Immunohistochemistry confirmed that CaMKHα^+^ excitatory neurons of the vHP (**Figure 1B**), but not GABAergic inhibitory neurons, expressed ChR2-mCherry (**Supplementary Figure 1A**).

**Figure 1.**
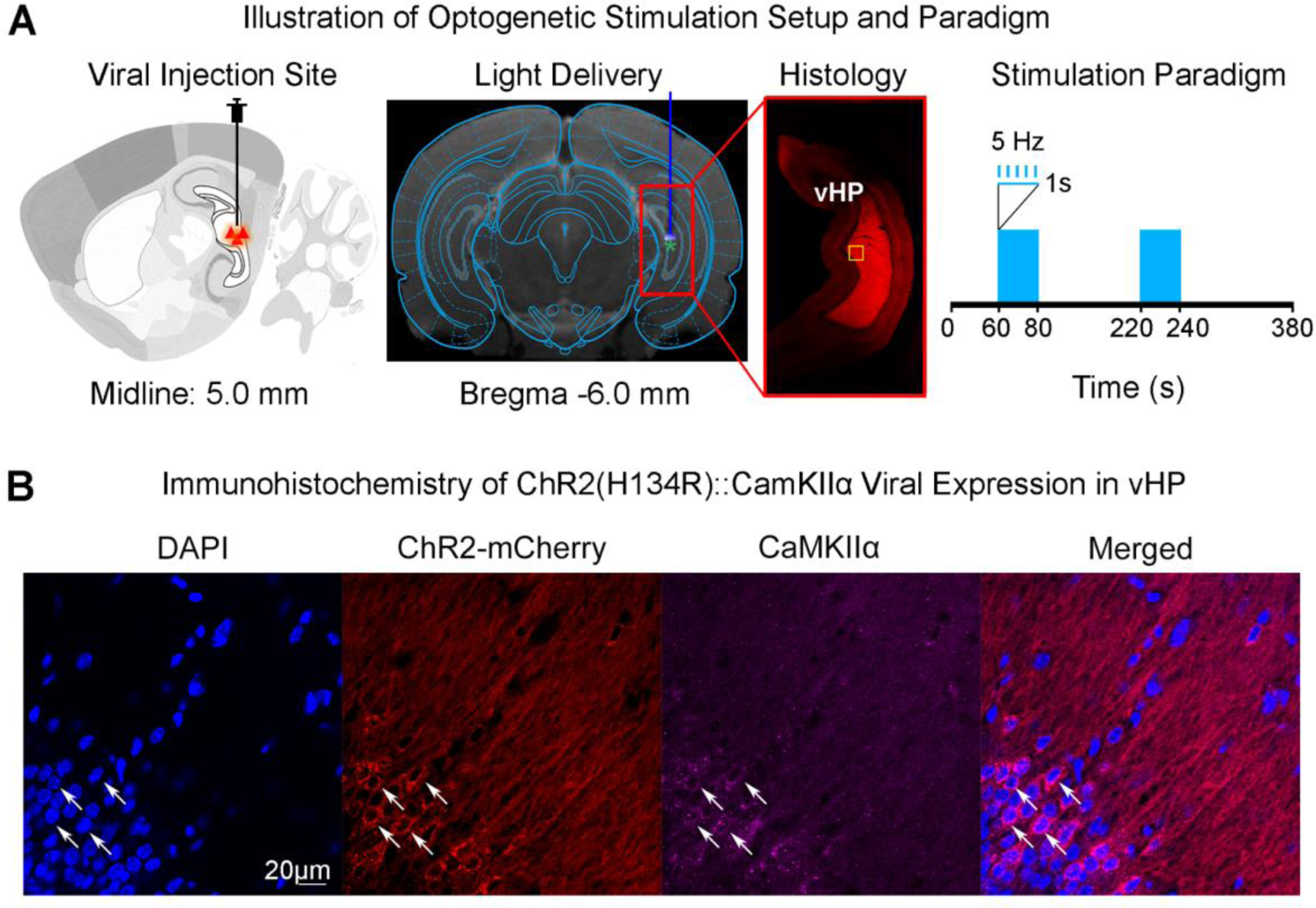
Experimental setup for optogenetic stimulation and histological characterization of ChR2::CaMKII viral expression in ventral hippocampus (vHP) excitatory neurons. *(A)* Schematic *(Left)* and T2-weighted anatomical image *(Middle)* shows the viral injection and fiber implantation sites, respectively. Histology image shows viral expression in vHP (Red box). The yellow box indicates the location of magnified confocal images shown in *B*. Optogenetic fMRI stimulation paradigm *(Right)*. 5 Hz was presented at 30 % duty cycle in a block-design paradigm (20 seconds ON; 140 seconds OFF). *(B)* Merged representative confocal images co-stained for the nuclear marker DAPI, ChR2-mCherry, and excitatory marker CaMKIIα^+^ neurons of vHP (white arrows).

To examine frequency-dependent spatiotemporal characteristics of brain-wide, long-range evoked BOLD responses driven by vHP, we performed whole-brain optogenetic fMRI in lightly anesthetized rats. Blue light pulses at five frequencies (1 Hz with 10 % duty cycle, 5 Hz, 10 Hz, 20 Hz, and 40 Hz with 30 % duty cycle; light intensity, 40 mW/mm^2^) were delivered to vHP neurons in a block design paradigm (**Supplementary Figure 2A**). We chose a reduced duty cycle for 1 Hz stimulation to avoid excessively long stimulation pulse width that may not be physiological. 5 Hz optogenetic stimulation of vHP evoked robust brain-wide positive BOLD activations in regions related to learning, memory, sensory processing and emotion, including bilateral vHP, dorsal hippocampus (dHP), entorhinal cortex (Ent), primary auditory cortex (AC), perirhinal cortex (Prh), amygdala (AMG), medial septum (MS), lateral septum (LS), diagonal band of Broca (DBB), and cingulate cortex (Cg) (**Figure 2**). Note that we did not detect any positive BOLD activations in IC and MGB in midbrain and thalamus, respectively. Such brain-wide activations evoked by 5 Hz stimulation indicated a high possibility of sound processing modulation. Further, this frequency matches the previously reported range of hippocampal theta oscillations, which travels along the hippocampal septotemporal axis (44, 45). Importantly, we found robust BOLD activations in the AC only during 5 Hz stimulation at vHP, but not at the other four frequencies (1 Hz, 10 Hz, 20 Hz, 40 Hz) (**Supplementary Figure 2C**). These frequencies evoked weaker BOLD responses in Prh, MS, and LS, and Cg, while retaining strong BOLD responses in vHP and dHP. Altogether, these results demonstrate the most extensive brain-wide vHP downstream targets found at 5 Hz stimulation, especially the robust BOLD activations in AC. These findings indicate that 5 Hz vHP stimulation generates strong and robust hippocampal activity outputs to AC and other regions, likely modulating sound processing brain-wide.

**Figure 2.**
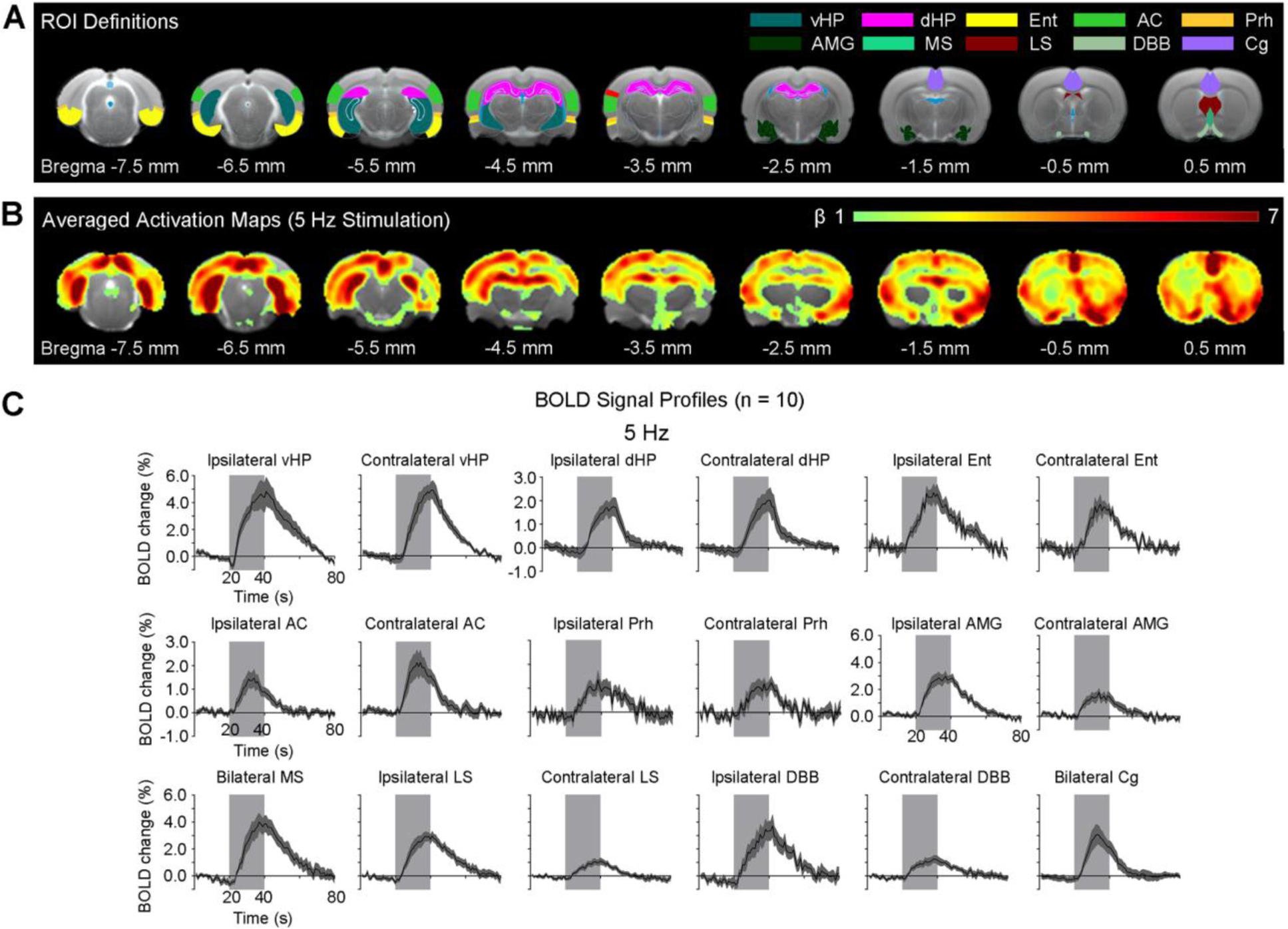
Brain-wide activations detected in the hippocampal formation, and cortical and subcortical regions during 5 Hz optogenetic stimulation of excitatory neurons in vHP. *(A)* Regions of interest (ROIs) defined by the rat brain atlas used to extract the BOLD signal profiles. *(B)* Averaged activation (β) maps of optogenetic stimulation in vHP. Robust positive BOLD responses detected in bilateral vHP, dHP, Ent, AC, Prh, AMG, MS, LS, DBB, and Cg during 5 Hz optogenetic stimulation (n = 10; t > 3.1; corresponding to p < 0.001). *(C)* BOLD signal profiles extracted from the ROIs. Error bars indicate ± SEM. The area shaded in grey indicates the 20 s 5 Hz optogenetic stimulation window. Abbreviations: Ventral Hippocampus (vHP); Dorsal Hippocampus (dHP); Entorhinal Cortex (Ent); Auditory Cortex (AC); Perirhinal Cortex (Prh); Amygdala (AMG); Medial Septum (MS); Lateral Septum (LS); Diagonal Band of Broca (DBB); Cingulate Cortex (Cg).

### Hippocampal outputs enhance neural responses and their selectivity to vocalizations with negative valence in auditory midbrain, thalamus and cortex

To explore the large-scale hippocampal modulatory effects on early auditory processing of natural sound, we performed auditory fMRI with and without continuous 5 Hz optogenetic stimulation at vHP. Forward aversive vocalizations (i.e., natural and behaviorally relevant) and the same but temporally reversed vocalizations (i.e., artificial and behaviorally irrelevant) were presented to the contralateral/left ear in a block-design paradigm (**Supplementary Figure 3A**). As expected, auditory evoked BOLD responses occurred along the ascending auditory pathway, including ipsilateral IC, MGB, and bilateral AC (**Figure 3**). Without optogenetic stimulation, the BOLD responses (as described by β values) in IC, MGB and AC were stronger using forward than reversed vocalizations (with β percentage difference between forward and reversed vocalization responses in IC: 8.6 ± 2.7 %, p < 0.01; MGB: 34.7 ± 10.53 %, p < 0.05; AC: 22.0 ± 7.7 %, p < 0.05, SDLUHG 6WXGHQW¶V t-test followed by Holm-Bonferroni correction). This finding demonstrates the response selectivity to forward vocalizations, which is consistent with our prior findings in rodent auditory system (46). Here, the response selectivity was most prominent in the external cortex (ECIC) and dorsal cortex (DCIC) of IC (with β percentage difference in ECIC: 8.8 ± 2.6 %, p < 0.01; DCIC: 7.8 ± 1.6 %, p < 0.01, SDLUHG 6WXGHQW¶V t-test followed by Holm-Bonferroni correction).

**Figure 3.**
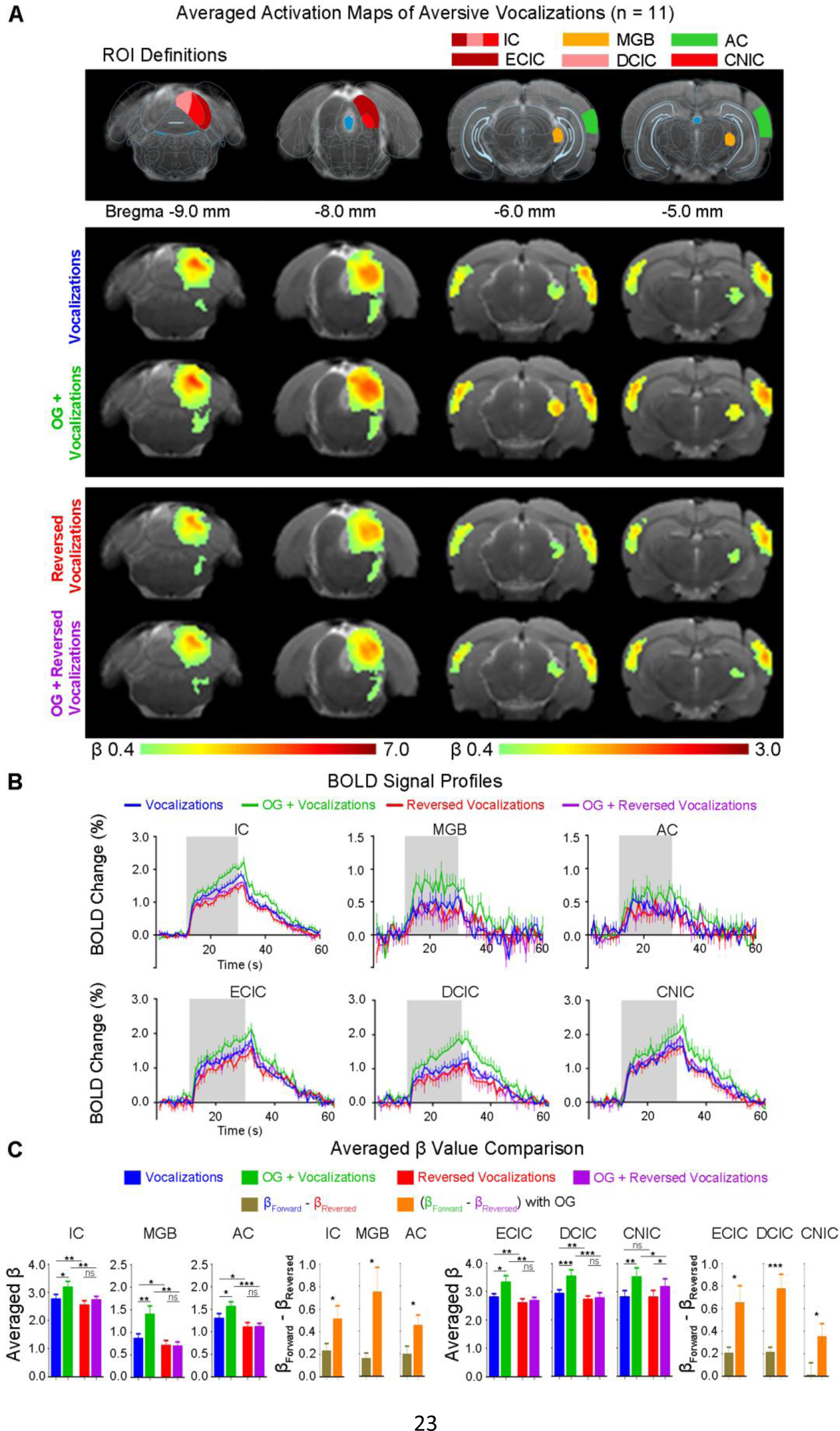
vHP optogenetic stimulation enhances neural responses and their selectivity to aversive vocalizations in the auditory midbrain (inferior colliculus or IC), thalamus (medial geniculate body or MGB), and cortex (auditory cortex or AC). *(A)* Illustration of the atlas-based region of interest (ROI) definitions *(Top).* Averaged BOLD activation (β) maps with and without 5-Hz optogenetic stimulation generated by fitting a canonical hemodynamic response function (HRF) to individual voxels in the fMRI image (n = 11; t > 2.6; corresponding to p < 0.01) *(Bottom)*. *(B)* BOLD signal profiles extracted from the corresponding ROIs (IC, MGB, AC, ECIC, DCIC, and CNIC). Error bars indicate ± SEM. The area shaded in grey indicates the 20 s acoustic stimulation. *(C)* BOLD signal (aYHUDJHG ȕ) comparison showing the modulatory effects of optogenetic stimulation on responses to forward aversive vocalizations in IC, MGB, AC, ECIC, DCIC, and CNIC, but not temporally reversed counterparts (that are artificial and evoke no behavioral response). Statistical comparisons were performed using paired two-sample t-test followed by Holm-Bonferroni correction with * for p < 0.05, ** for p < 0.01, *** for p < 0.001, and n.s. for not significant.

Optogenetic 5 Hz vHP stimulation significantly enhanced response selectivity to forward vocalizations throughout the ascending auditory pathway (with the β percentage difference between forward and reversed vocalization responses in IC: 16.3 ± 3.5 %, p < 0.01; MGB: 166.2 ± 57.6 %, p < 0.01; AC: 46.5 ± 11.9 %, p < 0.001, paired Student’s t-test followed by Holm-Bonferroni correction). Specifically, only the BOLD responses in IC, MGB and AC evoked by the forward vocalizations were significantly increased by optogenetic stimulation (with β percentage increase in forward vocalization response in IC: 18.2 ± 7.3 %, p < 0.05; MGB: 76.5 ± 22.6 %, p < 0.01; AC: 28.7 ± 14.1 %, p < 0.05, paired Student’s t-test followed by Holm-Bonferroni correction). This finding demonstrates that the hippocampal outputs selectively modulate the responses to forward vocalizations that convey contextual information. Note that such increased responses in IC occurred in ECIC and DCIC, as well as CNIC (with β increase in ECIC: 20.5 ± 9.5%, p < 0.05; DCIC: 20.8 ± 4.2 %, p < 0.001, CNIC: 26.3 ± 6.1 %, p < 0.01, paired Student’s t-test followed by Holm-Bonferroni correction). Altogether our fMRI results indicate that hippocampal outputs (initiated by the 5 Hz vHP stimulation) can enhance IC, MGB and AC auditory responses and their selectivity to natural and behaviorally relevant sounds at early processing centers within the ascending auditory pathway.

### Hippocampal outputs enhance neural responses and their selectivity to vocalizations with positive valence

We then utilized the same approach to examine whether such hippocampal modulation on natural sound processing was biased for only the aversive content in vocalizations. So, we performed auditory fMRI with postejaculatory vocalizations with and without presenting the 5 Hz optogenetic stimulation at vHP. Similarly, auditory evoked BOLD responses occurred along the ascending auditory pathway, including IC, MGB, and AC (**Figure 4**). Without optogenetic stimulation, the BOLD responses evoked by postejaculatory vocalizations also showed response selectivity to the forward one (with β percentage difference between forward and reversed vocalization responses in IC: 7.8 ± 1.7 %, p < 0.001; MGB: 43.0 ± 16.2 %, p < 0.05; AC: 40.5 ± 14.4 %, p < 0.05, paired Student’s t-test followed by Holm-Bonferroni correction). Consistent with the results of aversive vocalization experiment, the response selectivity to forward postejaculatory vocalizations in IC was mainly observed in ECIC and DCIC (with β percentage difference in ECIC: 5.4 ± 1.8 %, p < 0.01; DCIC: 6.5 ± 2.5 %, p < 0.05, paired Student’s t-test followed by Holm-Bonferroni correction), but not CNIC (no significant difference).

During optogenetic stimulation, similar to the aversive vocalization experiment, the response selectivity to the forward postejaculatory vocalizations was significantly enhanced throughout the ascending auditory pathway (with the β percentage difference between forward and reversed vocalization responses in IC: 19.2 ± 3.5 %, p < 0.001; MGB: 118.1 ± 43.0 %, p < 0.01; AC: 84.7 ± 23.2 %, p < 0.001, paired Student’s t-test followed by Holm-Bonferroni correction). Specifically, such enhancement primarily arose from increased responses to forward vocalizations (with β percentage increase in forward vocalization response in IC: 15.2 ± 4.8 %, p < 0.01, MGB: 123.1 ± 36.6%, p < 0.01; AC: 74.7 ± 34.2 %, p < 0.05, paired Student’s t-test followed by Holm-Bonferroni correction). In IC, such increased responses were found in ECIC, DCIC as well as CNIC (with β increase in ECIC: 16.0 ± 6.3 %, p < 0.05; DCIC: 9.8 ± 3.0 %, p < 0.01; CNIC: 17.3 ± 4.3 %, p < 0.01, paired Student’s t-test followed by Holm-Bonferroni correction). Taken together, these results demonstrate that the hippocampus plays a key role in modulating natural vocalization processing. Such hippocampal modulatory effects are not biased towards the aversive content embedded in the vocalizations. These fMRI findings again show that, for behaviorally relevant sound, the hippocampal outputs triggered by the 5 Hz vHP stimulation enhance auditory processing at large scale throughout the early ascending auditory pathway.

### Hippocampal outputs do not modulate neural responses to broadband acoustic noise

To further investigate whether the modulatory effects of hippocampal outputs are only specific to auditory processing of natural sounds, we replaced the vocalizations with a basic acoustic stimulus, 1 ± 40 kHz broadband noise (**Supplementary Figure 4**). As expected, the broadband noise evoked BOLD responses along the ascending auditory pathway, including IC, MGB, and AC. Importantly, the noise evoked BOLD responses in IC, MGB, and AC remained unchanged during the vHP stimulation (**Figure 5**). This finding reveals that the hippocampal outputs do not modulate behaviorally irrelevant or basic acoustic stimuli.

**Figure 4.**
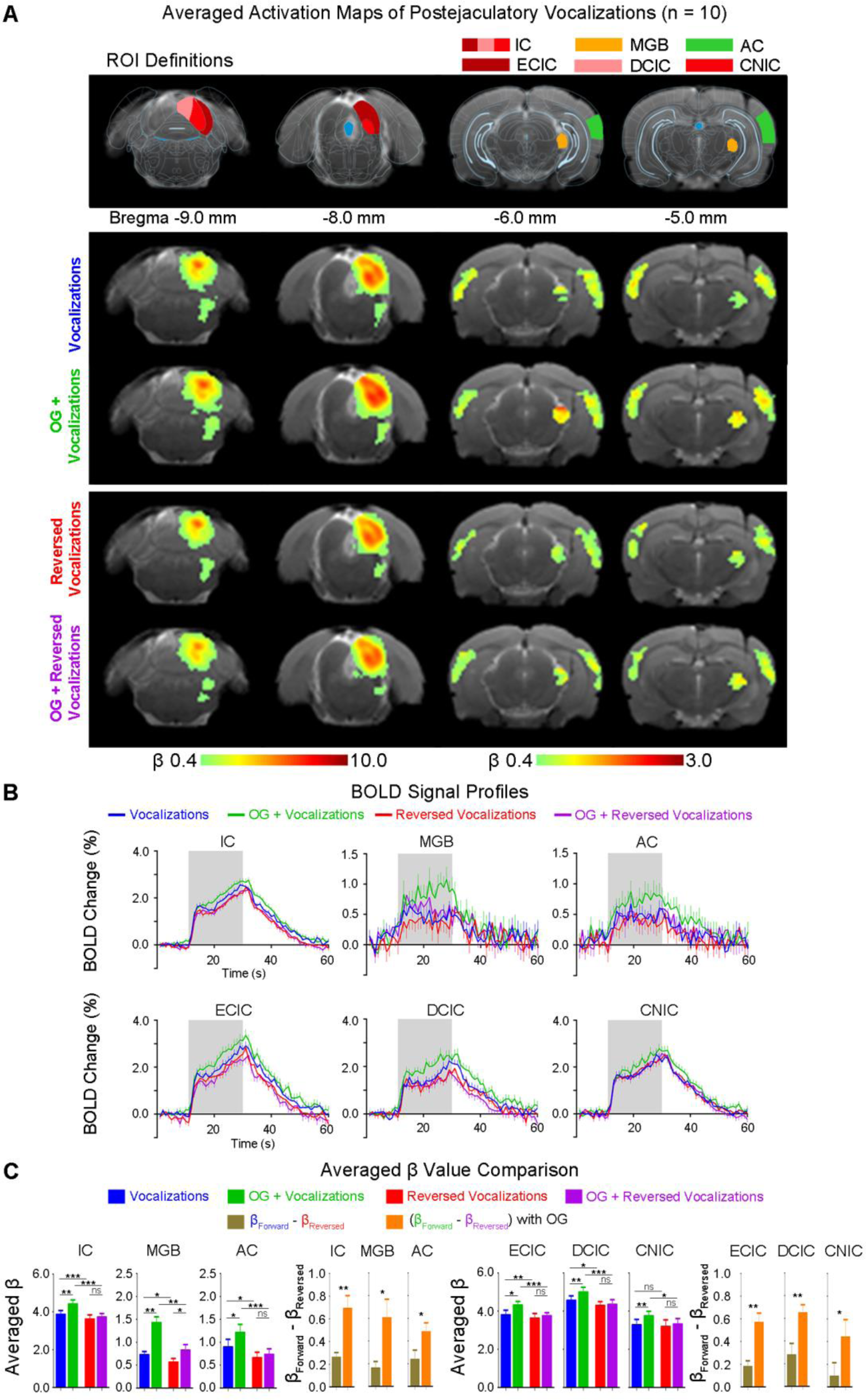
vHP optogenetic stimulation enhances neural responses and their selectivity to postejaculatory vocalizations in the auditory midbrain (IC), thalamus (MGB), and cortex (AC). *(A)* Illustration of the atlas-based region of interest (ROI) definitions *(Top)*. Averaged BOLD activation (β) maps with and without 5-Hz optogenetic stimulation *(Bottom)* generated by fitting a canonical HRF to individual voxels in the fMRI image (n = 10; t > 2.6; corresponding to p < 0.01). *(B)* BOLD signal profiles extracted from the corresponding ROIs (IC, MGB, AC, ECIC, DCIC, and CNIC). Error bars indicate ± SEM. The area shaded in grey indicates 20 s acoustic stimulation. *(C)* BOLD signal (aYHUDJHG ȕ) comparison showing the modulatory effects of optogenetic stimulation on responses to forward postejaculatory vocalizations in IC, MGB, AC, ECIC, DCIC, and CNIC, but not the temporally reversed counterparts. Statistical comparisons were performed using paired two-sample t-test followed by Holm-Bonferroni correction with * for p < 0.05, ** for p < 0.01, *** for p < 0.001, and n.s. for not significant.

**Figure 5.**
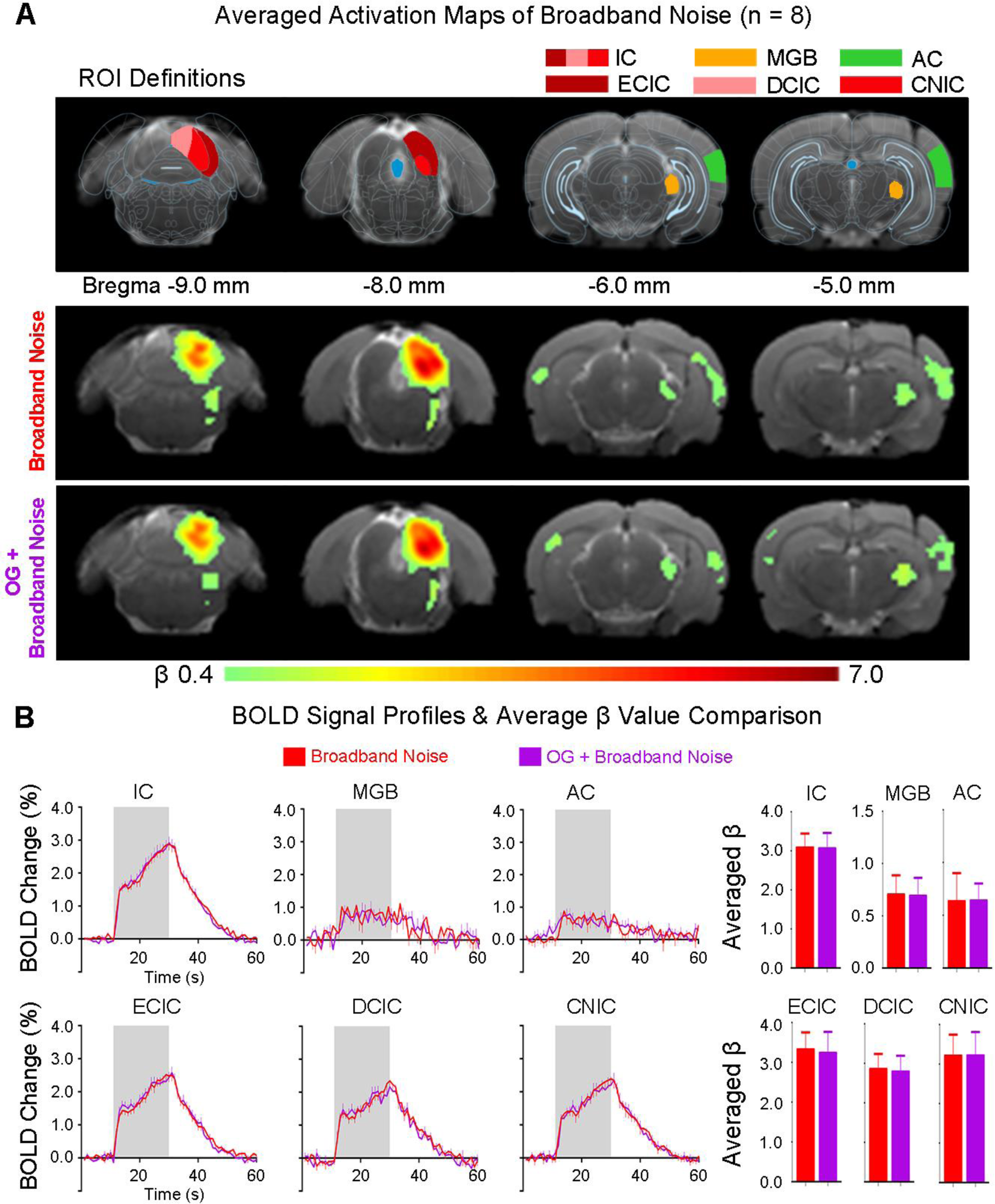
vHP optogenetic stimulation shows no modulatory effects on the responses to broadband ic noise in the auditory midbrain (IC), thalamus (MGB), and cortex (AC). *(A)* Illustration of the atlas-based region of interest (ROI) definitions *(Top)*. Averaged BOLD activation (β) maps with, without 5-Hz optogenetic stimulation *(Bottom)* generated by fitting a canonical HRF to individual voxels in the fMRI image (n = 8; t > 2.6; corresponding to p < 0.01). ***(B)*** BOLD signal profiles *(Left)* extracted from the corresponding ROIs (IC, MGB, AC, ECIC, DCIC, and CNIC). Error bars indicate ± SEM. The area shaded in grey indicates the 20 s acoustic stimulation. BOLD signal (aYHUDJHG ȕ) comparison *(Right)* showing no effects of optogenetic stimulation on broadband noise responses in IC, MGB, and AC. Statistical comparisons were performed using paired two-sample t-test followed by Holm-Bonferroni correction.

### Pharmacological hippocampal inactivation alters auditory responses and their selectivity for vocalizations

In addition, we examined the effects of pharmacologically inactivating neurons in the dentate gyrus of vHP on natural sound processing using tetrodotoxin (TTX). Auditory fMRI was performed before (PRE) and after (POST) infusion of TTX at vHP (**Supplementary Figure 5**). As expected, before the TTX infusion, the BOLD responses evoked by aversive vocalizations showed response selectivity to the forward one (**Figure 6**) (with β percentage difference between forward and reversed vocalization responses in IC: 15.6 ± 1.9 %, p < 0.01; MGB: 77.6 ± 23.5 %, p < 0.05; AC: 132.2 ± 60.5 %, p < 0.05, paired Student’s t-test followed by Holm-Bonferroni correction). Similarly, the response selectivity to forward aversive vocalizations in IC was mainly observed in ECIC and DCIC (with β percentage difference in ECIC: 11.2 ± 4.1 %, p < 0.05; DCIC: 31.4 ± 8.8 %, p < 0.05, paired Student’s t-test followed by Holm-Bonferroni correction), but not in CNIC.

Notably, pharmacological vHP inactivation via TTX infusion abolished the response selectivity to forward aversive vocalizations throughout the ascending auditory pathway, including IC, MGB, and AC. In general, the BOLD responses to forward and reversed vocalizations were also reduced. Yet the BOLD responses to forward vocalizations were diminished by a much greater extent (with β percentage decrease in forward vocalization response in IC: -20.6 ± 3.9 %, p < 0.01; MGB: -54.4 ± 21.5 %, p < 0.05; AC: -44.9 ± 17.0 %, p < 0.05, paired Student’s t-test followed by Holm-Bonferroni correction). In IC, such decreased responses were mainly found in ECIC and DCIC (with β percentage decrease in ECIC: -12.9 ± 3.9 %, p < 0.05; DCIC: -18.3 ± 3.6 %, p < 0.01, paired Student’s t-test followed by Holm-Bonferroni correction), but not in CNIC. These findings present additional evidence that vHP modulates and shapes IC, MGB and AC response selectivity to behaviorally relevant sounds at early sound processing centers within the ascending auditory pathway.

## Discussion

Here, we experimentally revealed the large-scale modulatory effects of ventral hippocampal outputs on early sound processing within the ascending auditory pathway by monitoring auditory responses during optogenetic vHP stimulation or pharmacological inactivation using large-view fMRI. We discovered a robust hippocampal influence on BOLD responses to vocalizations, but not artificial/basic acoustic stimuli, in the auditory midbrain, thalamus, and cortex. These fMRI results directly support the large-scale and faciliatory influence of the hippocampus on natural sound processing in early auditory centers within the ascending auditory pathway.

### Pathways subserving hippocampal top-down modulation of natural sound processing in early auditory centers

In the classical view, the auditory cortex processes complex auditory features and provides corticofugal feedback to modulate the responses in IC and MGB (4, 19–22). In particular, the existing hierarchical notion of cortical processing (48–50) postulates that the AC decodes the spectrotemporal dynamic features of auditory inputs, which facilitate responses to complex natural acoustic stimuli like vocalizations (5). However, studies suggest that complex sound processing may also occur at early auditory centers (51, 52). Converging evidence indicates that extraction and processing of spectral and temporal features begin at the midbrain level for discriminating natural sounds (53, 54), such as vocalizations (46, 55, 56). Moreover, an electrophysiological study showed that IC and MGB represent more comprehensive stimulus identities of natural stimuli relative to the cortex (4), highlighting the importance of early auditory structures in processing natural sounds. In addition, recent discoveries of non-canonical regions in processing complex auditory stimuli, such as entorhinal cortex (Ent) and medial septum (MS) (19, 20), and inevitably the hippocampus due to the dense reciprocal axonal projections (39, 57), challenge current dogma on the hierarchical notion of cortical processing. Here, we directly demonstrated that the hippocampus, a limbic region vital for learning and memory functions, modulates auditory responses to natural sounds along the early ascending auditory pathway. We propose that the hippocampus acts as a network hub that receives extensive projections from both subcortical and cortical regions (38, 58). In this view, multiple complementary pathways likely subserve long-range hippocampal modulation of central auditory processing of natural sound processing (**Figure 7**).

At the midbrain level, the IC integrates ascending auditory inputs from the lower auditory structures and descending feedback signals from the auditory thalamus and cortex (3). Despite the descending feedback from AC (24, 25), the IC also receives direct inputs from non-auditory structures such as the amygdala (AMG) (29), a limbic region that interacts with the hippocampus to regulate emotional memory (59, 60). The direct projection from AMG to IC may provide rapid feedback for processing emotional auditory stimuli (61). Such AMG-IC projections provide a route for vHP to modulate the responsiveness of IC neurons to specific types of stimuli based on the behavioral relevance and emotional valence (29, 61). Meanwhile, a recent retrograde tracing study revealed that IC receives descending projections from non-auditory regions, such as the entorhinal (Ent), perirhinal (Prh), and cingulate (Cg) cortices (28). These regions are part of the hippocampal formation and are linked with learning, memory, and sensory processes (31, 38, 62, 63). For instance, the hippocampal-entorhinal circuit can acquire and discriminate frequency and temporal features of auditory cues associated with reward-related behavior (20). Meanwhile, Prh inputs are required for fear conditioning to complex stimuli, such as vocalizations, but not for continuous tones (64, 65). Furthermore, the hippocampus also interacts with Cg to support memory encoding and retrieval of context-dependent information, thus facilitating the corresponding processing of behaviorally relevant stimuli (62, 63). Meanwhile, we found that our identified regions (AMG, Ent, Prh and Cg) were prominently activated upon 5 Hz optogenetic stimulation (**Figure 2**). Together, multiple afferents from non-auditory regions to IC suggest that inputs from higher-order structures, such as the vHP and its associated hippocampal formation, are vital for natural sound processing at the midbrain level.

At the thalamic level in the auditory hierarchy (66, 67), MGB can also receive hippocampal modulatory outputs via other sensory and prefrontal cortices, including somatosensory and cingulate cortices. For example, hind paw stimulation provides somatosensory inputs that can modulate auditory responses in the MGB (68), and electrical stimulation at the prefrontal cortex can influence spontaneous firing in the MGB (71). Meanwhile, MGB can also receive indirect inputs from the AMG via the thalamic reticular nucleus (TRN), which is critical for deviant sound detection (72). Exciting AMG-TRN projections amplified the sound-evoked responses in the auditory thalamus and, in turn, the cortex (73). Inactivating the basal AMG reduced certain conditioned responses to sound in MGB (74). Hence, the amplified sound-evoked response in MGB observed here can undergo modulation by hippocampal outputs via AMG and the prefrontal cortices such as cingulate.

At the cortical level, the auditory association area can receive hippocampal outputs directly from ventral CA1 neurons (41) and indirectly via the Ent, Prh, and parahippocampal cortices (75). Previous works have identified the functional roles of these hippocampal-cortical pathways comprising auditory recognition (20, 76, 77) and auditory-related memory processes (33, 36). Hippocampal modulatory outputs could potentially reach the auditory cortex through hippocampal-cortical pathways (37), and subsequently also modulate sound responses in the midbrain and thalamus via corticofugal projections (7, 23).

Recent evidence indicates that the reticular limbic auditory pathway may provide a fast route to relay auditory inputs from the cochlear nucleus to high-order regions (19), suggesting that the hippocampus receives and processes behaviorally relevant auditory inputs via Ent and MS. Further, our histological findings showed strong anterograde ChR2 mCherry expression in the MS and lateral septum (LS), as well as the diagonal band of Broca (DBB) (**Supplementary Figure 1B**), suggesting a circuit loop that is dedicated for auditory processing outside of the central pathways. These regions support learning (78) and memory functions (79), particularly related to auditory processes. For instance, MS inactivation impairs acquiring auditory fear memory (22). MS and DBB are the primary sources of cholinergic projections to HP and AMG (80), which can be critical for contextual memory formation (81), sensory cue detection and discrimination (82, 83). Disrupting MS and DBB cholinergic projections to vHP prevents auditory fear memory acquisition and retention (84). Further, a prior study showed that systemic blockade of cholinergic signaling via atropine, a muscarinic acetylcholine receptor antagonist, inhibited response selectivity to vocalizations in the auditory midbrain (46). Here, robust activations in the septum complex (i.e., MS, LS, and DBB) initiated from vHP (**Figure 2**) may trigger rapid downstream signaling cascades in the cholinergic system (82) and evoke postsynaptic responses at the terminal fields (85), such as the AMG. Cholinergic signaling within AMG is crucial for encoding and processing emotionally salient memories (86), which could facilitate selective amplification of auditory responses evoked by behaviorally relevant stimuli. Overall, we detected robust activations in cortical (i.e., AC, Ent, Prh, Cg) and subcortical regions (i.e., AMG, MS, LS, DBB) during 5 Hz optogenetic stimulation at vHP (**Figure 2**). The hippocampal modulatory outputs evoked from vHP could propagate and functionally interact with these activated regions, thereby modulating auditory responses at early auditory centers within the ascending auditory pathway, particularly at the midbrain and thalamic levels.

### Hippocampal outputs enhance natural sound responses at early auditory centers

In this study, we revealed that optogenetic activation of vHP, a region often associated with motivation or emotional behaviors (42, 43), enhanced auditory responses in early ascending auditory processing centers (i.e., IC, MGB) and the AC to forward, but not temporally reversed, vocalizations. Pharmacological inactivation of vHP diminished the auditory responses to forward vocalizations. Further, such hippocampal modulatory effects were absent when processing broadband noise, confirming that hippocampal activity is an integral component of natural sound processing within the ascending auditory pathway. Our findings demonstrate that the hippocampus is key for processing vocalizations and shaping the corresponding response selectivity.

The exact mechanisms underlying our experimentally observed selective modulation of natural sound processing by optogenetically triggered hippocampal outputs requires further investigation. We posit that specific spectrotemporal information embedded in the sound drive this specificity. The hippocampus is positioned to process temporal information of sensory inputs (33, 93), as hippocampal lesions can impair memory for the temporal order of events in both animals (94, 95) and humans (96, 97). When learning to associate specific time intervals with a given stimulus, the hippocampus is essential for discriminating minute temporal differences in rodents (98). These findings indicate that the hippocampus plays a critical role in encoding and recognizing temporal information to subsequently discriminate and interpret the temporal organization of incoming sensory inputs.

Here, we observed that hippocampal modulation primarily facilitates auditory responses in IC, MGB, and AC to both types of vocalizations (i.e., aversive/fear and postejaculatory/positive), but not their temporally reversed counterparts (**Figures 3** and **4**). We postulate that temporally reversing vocalizations alter specific properties. Reversed vocalizations no longer carry the critical information embedded in forward vocalizations, which diminishes the behavioral relevance of the sound. The hippocampus, by interpreting and discriminating the embedded spectrotemporal features of the incoming sounds, can discriminate and recognize natural and/or behaviorally relevant stimuli by contextual memory recall according to past experience (99, 100) and then can exert selective influence to downstream targets. Moreover, blocking hippocampal outputs through pharmacological manipulation altered responses to forward vocalizations, and consequently disrupted the response selectivity to vocalizations (**Figure 6**). Together, our results showed that hippocampal outputs are critical to facilitate auditory responses to natural sounds.

**Figure 6.**
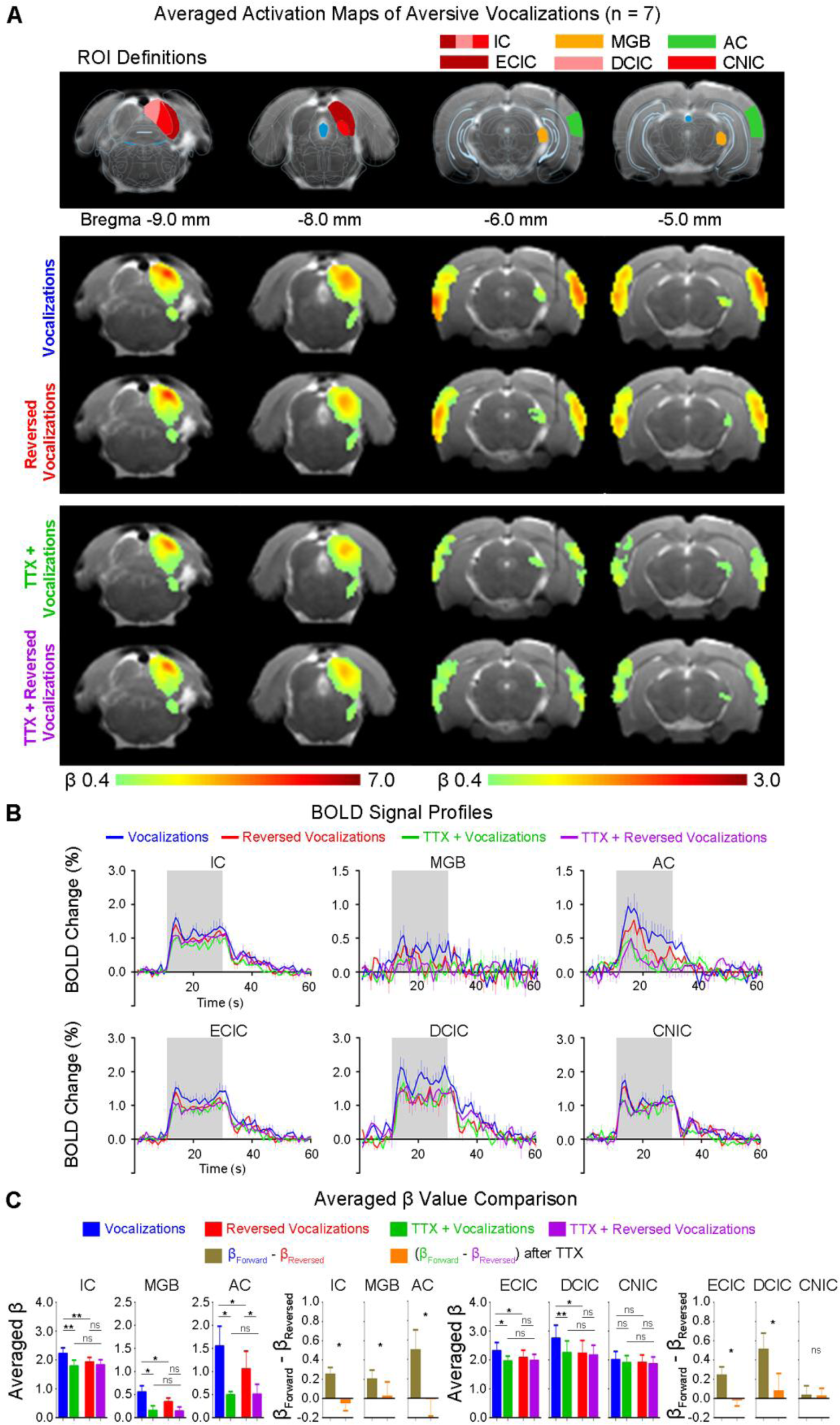
Pharmacologically inactivating vHP alters neural responses and decreases their selectivity to aversive vocalizations in the auditory midbrain (IC), thalamus (MGB), and cortex (AC). *(A)* Illustration of the atlas-based region of interest (ROI) definition *(Top)*. Averaged BOLD activation (β) maps before and after TTX infusion *(Bottom)* generated by fitting a canonical hemodynamic response function (HRF) to individual voxels in the fMRI image (n = 7; t > 2.6; corresponding to p < 0.01). ***(B)*** BOLD signal profiles extracted from the corresponding ROIs (IC, MGB, AC, ECIC, DCIC, and CNIC). Error bars indicate ± SEM. The area shaded in grey indicates 20 s acoustic stimulation. ***(C)*** BOLD signal (averaged β) comparison showing the effects of TTX inactivation of vHP neurons predominantly on responses to forward aversive vocalizations in IC, MGB, AC, ECIC, and DCIC. Statistical comparisons were performed using paired two-sample t-test followed by Holm-Bonferroni correction with * for p < 0.05, ** for p < 0.01, *** for p < 0.001, and n.s. for not significant.

**Figure 7.**
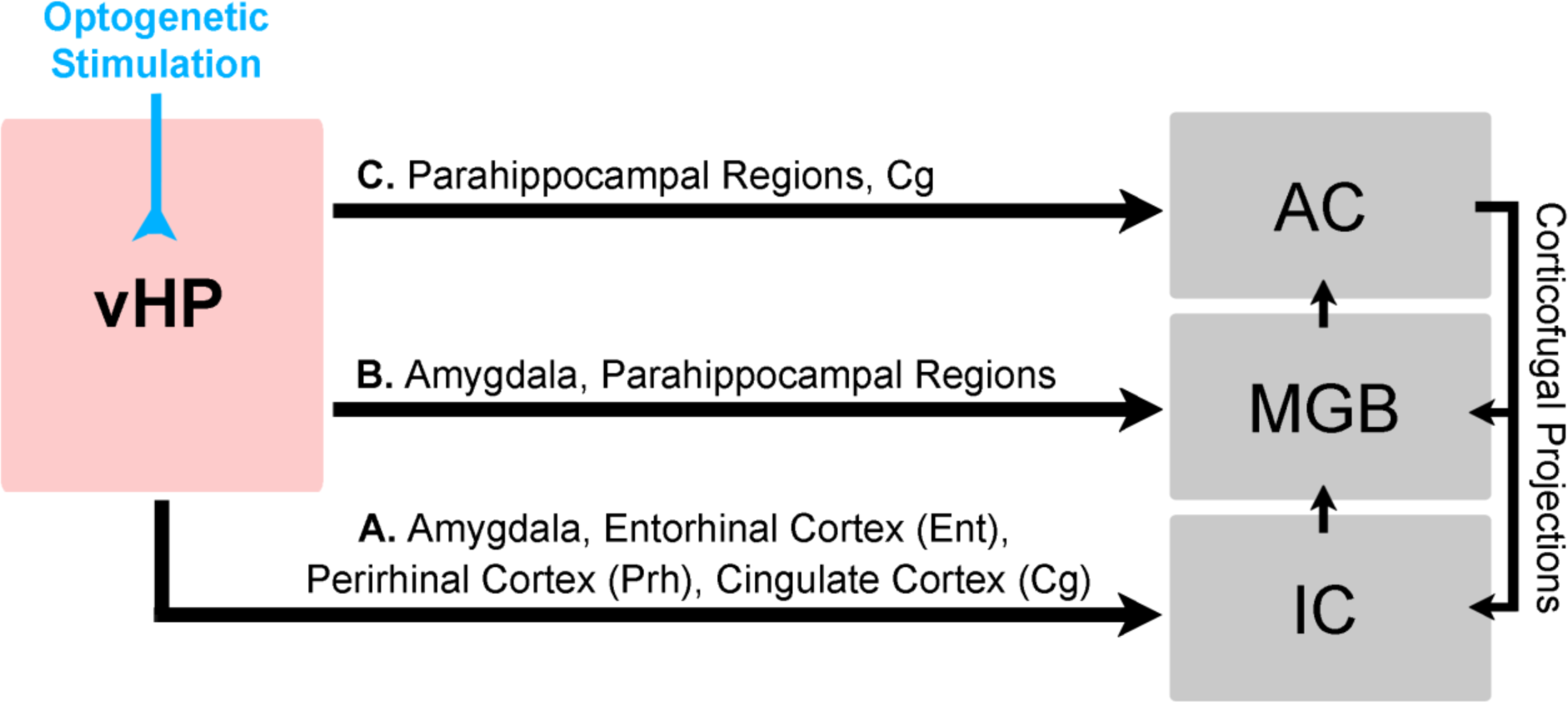
Schematic pathways of long-range hippocampal modulation of natural sound processing within the ascending auditory pathway through optogenetically-evoked vHP activity. *Route A:* vHP could modulate responses in IC via indirect projections from amygdala, entorhinal cortex (Ent), perirhinal cortex (Prh) and cingulate cortex (Cg), which can then enhance the responses in MGB and AC along the ascending auditory pathway. *Route B:* vHP can modulate MGB responses indirectly via amygdala and parahippocampal regions, which could subsequently modulate responses in AC via ascending auditory pathway. *Route C:* vHP could modulate AC responses directly via hippocampal-cortical projections or indirectly via parahippocampal regions such as Ent, Prh, and Cg via cortico-cortical projections. AC could then modulate the responses in MGB and IC via corticofugal projections.

Hippocampal modulatory outputs, such as the increased coherence of neuronal firing with theta/gamma oscillations (101) and/or modifying synaptic strengths (102), can promote communication between the hippocampus and its downstream targets, and then facilitate natural sound processing via multiple pathways. At a systems level, theta oscillations can correspond to contextual memory recall (i.e., retrieval of contextual information pertaining to a specific event in the past where its temporal context or sequence is important) (103–105). During memory retrieval, theta oscillations coupled with gamma oscillations were enhanced when processing previously learned, behaviorally relevant stimuli (106). We speculate that optogenetically initiated hippocampal outputs generate theta-like oscillations in the hippocampal-cortical network (44, 45) to enhance retrieval processes between the hippocampus and Ent during auditory processing. Previous studies also demonstrated the importance of hippocampal and cortical theta oscillations (101, 107) in coordinating groups of neurons to integrate sensory information (108). Theta oscillations are essential to coordinate brain-wide activity (109), such that neuronal spiking in somatosensory (110), prefrontal (111) and entorhinal cortices (112) are phase-locked to hippocampal theta oscillations. These studies indicate that the hippocampus can influence distal sensory responses via theta oscillations.

In this study, we uncovered long-range hippocampal modulation of natural sound processing within the ascending auditory pathway during 5 Hz optogenetic stimulation at vHP. This stimulation frequency coincides with the reported range of hippocampal theta oscillations (101, 107), evoked the most robust brain-wide activations, including in primary AC. Such theta-like activities could trigger the propagation of hippocampal activity to distal regions, facilitating the interaction with neural activity in the ascending auditory pathway (i.e., AC, MGB and IC) and other targets to produce modulatory effects. However, we do not preclude the potential modulatory effects of other stimulation frequencies in the range of theta oscillations like 10 Hz (101, 107). Even though BOLD activations evoked by 10 Hz optogenetic stimulation were not as widespread as 5 Hz, we still observed robust activations in subcortical regions (e.g., AMG, MS, LS, DBB) albeit with weaker activations in cortical regions (e.g., AC, Prh, Cg) (**Supplementary Figure 2**). Overall, our findings suggest that hippocampal top-down modulatory outputs, which may be triggered by behaviourally relevant auditory inputs, were augmented by optogenetic stimulation of vHP to enhance responses to natural sounds.

In summary, the present fMRI study established a top-down and large-scale modulatory role for the hippocampus throughout the ascending auditory pathway, including the auditory midbrain, thalamus and cortex, to faciliate natural sound processing. Our findings expand our present understanding of central auditory system beyond the traditional cortex centric views. Future studies should elucidate the precise hippocampal modulatory processes of natural sound that arise from the brain-wide auditory information processing networks.

## Methods

### Subjects

Adult male Sprague-Dawley rats were used in all experiments. Animals were individually housed under a 12-h light/dark cycle with access to food and water ad libitum. All animal experiments were approved by the Committee on the Use of Live Animals in Teaching and Research of the University of Hong Kong. Group I underwent optogenetic fMRI experiments (n = 10), group II underwent combined optogenetics and auditory fMRI experiments (n = 11, aversive vocalizations experiments; n = 10, postejaculatory vocalizations experiments; n = 8, broadband noise experiments), and group III underwent combined pharmacological and auditory fMRI experiments (n = 7, aversive vocalizations experiments). Full details of animal surgical procedures, optogenetic stimulation paradigms, combined optogenetics, and auditory fMRI acquisition and analysis procedures, and histology are provided in SI Methods.

### Data and Code Availability

The data files that support the findings of this study and computer codes used are available on Dryad Digital repository (https://doi.org/10.5061/dryad.08kprr52x).

## Acknowledgments

This work was supported by Hong Kong Research Grant Council (C7048-16G and HKU17112120 to E.X. Wu, and HKU17103819 and HKU17104020 to A.T.L. Leong), Lam Woo Foundation, Guangdong Key Technologies for Treatment of Brain Disorders (2018B030332001) and Guangdong Key Technologies for AO]KHLPHU¶V Disease Diagnostic and Treatment (2018B030336001) to E.X. Wu. We would like to thank Profs. J. He and G. Buzsáki for the insightful scientific discussions. We also thank Drs. R. Chan, C. Dong, A. To, and M. Bialy for their technical assistance. We also thank Dr. K. Deisseroth who provided us with the ChR2 viral construct.

## SI Appendix

### SI Methods

#### Virus Packaging

Channelrhodopsin2-mCherry fusion protein under the control of the Ca^2+^ /calmodulin- dependent protein kinase IIa (CaMKIIa) promoter was used. The AAV5-CaMKIIa- ChR2(H134R)-mCherry plasmid (maps available online from www.stanford.edu/group/dlab/optogenetics) was packaged by the viral vector core of the University of North Carolina at Chapel Hill, Chapel Hill, NC (tire of 4 x 10^12^ particles/mL).

#### Stereotactic Surgery for Viral Injection

Stereotactic surgery was performed when rats were 6 weeks old with bodyweight around 250 g. Rats were anesthetized with an intraperitoneal bolus injection of ketamine (90 mg/kg) and xylazine (40 mg/kg) mixture. The scalp was shaved, and the rats were secured in a stereotactic frame. Buprenorphine (0.05 mg/kg) was administered subcutaneously to minimize pain, and heating pads were used to prevent hypothermia. Following a midline incision, a craniotomy was made on the right hemisphere in the vHP, and injection was performed at two depths (-6.00 mm posterior to Bregma, +5.00 mm ML, -4.75 and -4.50 mm from brain surface). For optogenetic animals, 1.5 µL of viral constructs were delivered through a 5 µL syringe and 33-gauge beveled needle injected at 150 nL/min at each depth. Following viral injection, the needle was held in the place for 10 minutes before slow retraction. Then, the scalp incision was sutured. After the surgery, buprenorphine (0.05 mg/kg) was administered subcutaneously twice daily for 72 hours to minimize post-surgery infection and inflammation. Animals rested for six weeks before fMRI experiments were performed.

#### Optical Fiber Implantation

Stereotactic surgery was performed to implant custom made plastic optical fiber cannula (POF, core diameter 450 µm; Mitsubishi Super ESKA^TM^ CK-20) at the viral injection site 1 – 2 hours before fMRI experiments. Rats were anesthetized with isoflurane (induction 3 % and maintenance 2 %) and secured on a stereotactic frame. Following a midline incision, a craniotomy was made at the same coordinates as the viral injection site. A heating pad was used to prevent hypothermia. Before implantation, the fiber tip was beveled to facilitate insertion and minimize injury to brain tissue. Then, it was inserted with the fiber tip at a depth of 4.7 mm. The optical fiber was fixed on the skull with UV glue and dental cement, and the scalp incision was sutured. The fiber outside the brain was made opaque using heat-shrinkable sleeves to avoid undesired visual stimulation. After the surgery, buprenorphine (0.05 mg/kg) was administered subcutaneously to minimize discomfort.

#### Cannula Implantation for Tetrodotoxin (TTX) Infusion

Before animals were placed in the magnet, surgery was performed to implant an MRI-compatible cannula, 250-µm internal diameter in vHP. Before surgery, animals were anesthetized with isoflurane (induction 3% and maintenance 2%) secured on a stereotactic frame. Buprenorphine (0.05 mg/kg) was administered subcutaneously to minimize discomfort before MRI experiments. The concentration of TTX used was 5-10 ng/µL, similar to the values used in previous in vivo studies (113, 114).

#### Animal Preparation for MRI Experiments

All MRI experiments were carried out on a 7T MRI scanner (PharmaScan 70/16, Bruker Biospin) using a transmit-only birdcage coil in combination with an actively decoupled receive-only surface coil. After surgery, one to two drops of 2% lidocaine was applied to the chords to provide local anesthesia before endotracheal intubation. The animals were mechanically ventilated at a rate of 60 breaths per minute with 1–1.5% isoflurane in room-temperature air using a ventilator (TOPO, Kent Scientific). During all fMRI experiments, animals were placed on a plastic cradle with the head fixed with a tooth bar and plastic screws in the ear canals. Rectal temperature was maintained at ∼37.0 °C using a water circulation system. Continuous physiological monitoring was performed using an MRI-compatible system (SA Instruments). Vital signs were within normal physiological ranges (rectal temperature: 36.5 – 37.5 °C, heart rate: 350 – 420 beat/min, respiration rate: 60 breath/min, oxygen saturation: > 95%) throughout the experiments.

#### MRI-Synchronized Optogenetic and Auditory Stimulation

An Arduino programming board synchronized the scanner trigger and the lasers for optogenetic and visual stimulation. Computers and light delivery systems were kept outside the magnet, and long optical patch cables (5–10 m) delivered light into the bore of the scanner. For optogenetic stimulation, blue light was delivered using a 473-nm DPSS laser. Light intensity was measured (PM100D, Thorlabs, USA) before each experiment as 8 mW at the fiber tip (450 µm, NA = 0.5), corresponding to a light intensity of 40 mW/mm^2^. For auditory stimulation, acoustic stimuli were controlled by a computer and produced by a high frequency multi-field magnetic speaker (MF1, TDT) driven by an amplifier (SA1, TDT). Monaural stimulation was delivered through a custom- made 165 cm long rigid tube and a 6.5 cm soft tube into the animals’ left ear canal. The right ear was occluded with cotton and Vaseline, to reduce the scanner noise reaching the ears. The sound pressure level (SPL) was measured by a recorder (FR2, Fostex, Japan) placed at ∼ 2 mm from the tip of the soft tube. The variance of the light power was maintained less than 2.5 mW/mm^2^ and the SPL less than 2 dB. This setup has been used in our previous studies (46, 115).

To determine the frequency-dependent spatiotemporal characteristics of evoked vHP responses (optogenetic fMRI experiments), five frequencies were used (1 Hz, 5 Hz, 10 Hz, 20 Hz, and 40 Hz) with a light intensity of 40 mW/mm^2^. 30 % duty cycle was used for all stimulation frequencies, except 1 Hz, which was at 10 % duty cycle. The duty cycle for 1 Hz optogenetic stimulation was reduced to avoid a very long stimulation pulse width which may not be physiological. vHP excitatory neurons were stimulated with a block design paradigm that consisted of 60 seconds light-off followed by 20 seconds light-on and 140 seconds light-off periods. Three to four trials were recorded for each frequency in an interleaved manner in each animal.

In combined optogenetic and auditory fMRI experiments, the effects of optogenetic stimulation in the vHP on brain baseline BOLD signals were examined by presenting 5 Hz stimulation (60 seconds light-off followed by 20 seconds light-on and 140 seconds light-off periods), without presenting acoustic stimulation. This paradigm was repeated twice in each animal. Subsequently, the effects of optogenetic stimulation on auditory midbrain, thalamus, and cortex processing of sound stimulation were investigated.

In the vocalization experiment, two types of vocalizations (I. ‘Aversive’ Vocalizations: bandwidth: 22-25 kHz, peak frequency: 22 kHz; sound pressure level (SPL): 83 dB, obtained online from http://www.avisoft.com/rats.htm (46); II. ‘Postejaculatory’ Vocalizations: bandwidth: 22 kHz, peak frequency: 22 kHz; SPL: 83 dB) from (91) and their temporal reversions were presented. Standard block-design paradigm was used for the auditory stimulation (40 s sound-off followed by 4 blocks of 20 s sound-on and 40 s sound-off, fMRI no. of time points = 280). During every 20 s sound-on period, the aversive vocalization (length 1.2 s, plus silence 0.8 s afterward) was repeated ten times at 60 % duty cycle, whereas the postejaculatory vocalization (length 3.6 s, plus silence 0.4 s afterward) were repeated five times at 90 % duty cycle. For each animal, this paradigm was repeated six times for each type of vocalization sounds. For three of them, the forward vocalization was presented first; and for the other three, the reversed one was presented first. Note that the forward and temporally reversed vocalizations contained identical acoustic features except reversed temporal information (**Supplementary Figure 3**). Note that temporally reversed vocalizations were employed here as the control for forward (i.e., true and behaviorally relevant) vocalizations. Such temporally reversed vocalizations exhibited the identical sound pressure level (SPL that is important to BOLD response level), but triggered minimal behavioral responses (46).

In the control experiment, broadband noise (bandwidth: 1 – 40 kHz; SPL: 83 dB) was presented to the left ear canal of the animals in a standard block-design paradigm (40 seconds sound-off followed by four blocks of 20 seconds sound-on and 40 seconds sound-off, fMRI no. of time points = 280) (**Supplementary Figure 4**). During each 20 seconds sound-on period, the broadband noise was presented with amplitude modulation at 4 Hz and 80 % duty cycle. The optogenetic stimulation (light wavelength: 473 nm, intensity: 40 mW/mm^2^, pulse rate: 5 Hz, duty cycle: 30%) was continuously presented to the right vHP throughout the auditory fMRI sessions and alternated between sessions.

In combined TTX and auditory fMRI experiment, a total of sixteen auditory fMRI sessions were performed in each animal. After eight sessions, 5µL TTX (concentration: 5-10 ng/µL) (113, 114) was infused into vHP. The next immediate session was then acquired one minute after the TTX infusion. During auditory fMRI sessions, auditory stimuli were presented to the left ear canal of the animals. Aversive vocalization and its temporal reversion were presented in standard block design paradigm (40 s sound-off followed by 4 blocks of 20 s sound-on and 40 s sound-off, fMRI no. of time points = 280). Auditory fMRI sessions were interleaved (i.e., either starting with forward aversive vocalization or temporally reversed aversive vocalization; **Supplementary Figure 5**).

#### MRI Acquisitions

Scout images were first acquired to determine the coronal and sagittal planes of the brain. Twelve coronal slices with 1.0 mm thickness were positioned to cover the ascending auditory pathway with the 2^nd,^ and 3^rd^ slice covered the whole IC. T2-weighted images were acquired as anatomical reference using a Rapid Acquisition with Refocused Echoes (RARE) sequence (FOV = 32 x 32 mm^2^, data matrix = 256 x 256, RARE factor = 8, echo time (TE) = 36, repetition time (TR) = 4200 ms). All fMRI measurements were obtained using a multi-slice single-shot Gradient-Echo Echo-Planar-Imaging (GE-EPI) sequence (FOV = 32 x 32 mm^2^, data matrix = 64 x 64, flip angle = 56°, TE/TR = 20/1000 ms, temporal resolution = 1000ms). Note that the fiber outside was made opaque using heat-shrinkable sleeves to avoid unwanted visual stimulation.

#### fMRI Data Analysis

For each animal, the fMRI images from each animal were realigned to the mean image of the first fMRI session (SPM12, Wellcome Department of Imaging Neuroscience, University College London, UK). Images from each animal were co-registered to a custom-made brain template using affine transformation and Gaussian smoothing, with the criteria of maximizing normalized mutual information (SPM12). For optogenetic fMRI and auditory fMRI, data from repeated sessions were averaged, in-plane smoothed (FWHM = 1 pixel), and high-pass filtered (128 s), and then standard general linear model (GLM) was applied, to calculate the BOLD response coefficient (β) maps for each stimulus (SPM12). fMRI sessions suffering from motion artifacts (>0.125 mm voxel shifts detected by realignment) and sudden physiological changes (i.e., abrupt changes in respiration pattern, heart rate and oxygen saturation level) were discarded. Typically, in each animal, three fMRI sessions were averaged for vHP optogenetic stimulation only, whereas four sessions were averaged for combined auditory and vHP optogenetic stimulation. Finally, activated voxels were identified with Student’s t-test on the β values (p < 0.01).

Three regions-of-interest (ROIs) covering different IC subdivisions were defined using the Paxinos & Watson rat brain atlas. The ROI that covered the inferior colliculus (IC), medial geniculate nucleus (MGB), or auditory cortex (AC) was defined by identifying clusters of activated voxels (p < 0.05) that were restricted within the anatomical location of each region. Anatomical locations of IC, MGB, and AC, were determined using the atlas. In individual animals, the BOLD signal profiles for each ROI were first extracted and averaged across voxels, before they were separated into six blocks (each covering a period from 10 s before to 30 s after a sound-on period) and two blocks (each covering a period from 10 s before to 50 s after an optogenetic-ON period), respectively. They were then averaged again and normalized by the mean signal intensity of the first 10 s to calculate the percentage of BOLD signal change. Final averaging was then performed across animals to generate BOLD signal profiles.

In individual animals, β values were also extracted from each ROI and averaged across voxels. The final β value used for comparison between the BOLD responses to the sound stimulus with and without optogenetic stimulation of the vHP was computed by averaging. Further, the β value difference between forward and reversed vocalizations (βForward - βReversed), as a metric of response selectivity, was compared between with and without optogenetic stimulation. Note that the size of ROI for each IC subdivision was different, and this could influence the absolute SNR of the averaged BOLD responses.

#### Histology, Immunohistochemistry, and Confocal imaging

Upon completion of in vivo studies, animals were deeply anesthetized with pentobarbital and transcardially perfused with ice-cold 4% paraformaldehyde (PFA) in PBS. Brains were extracted and fixed in 4% PFA for 4 h at 4 °C. The brains were equilibrated in 20% sucrose in PBS at 4 °C overnight. Axial sections (40 µm) were prepared on a freezing microtome (model 860, AO Scientific Instruments). Consecutive sections (120 µm apart) were mounted and examined with a laser confocal microscope (Carl Zeiss LSM780). For immunohistochemistry, free-floating sections were processed with 5% normal goat serum and 0.3% Triton X-100 in PBS with primary antibodies against rabbit polyclonal to CaMKIIa (1:400; Abcam) and guinea pig polyclonal to GABA (1:200; Abcam) at 4 °C for 24 h. After washing with PBS, sections were then incubated for 2 h at room temperature with secondary antibodies Alexa Fluor 647 conjugate goat anti-rabbit IgG and Alexa Fluor 488 conjugate goat anti-guinea pig IgG (both 1:500; Molecular Probe). Slices were then washed and mounted using FluoroShield mounting medium with DAPI (Abcam). Double or triple immunofluorescence was assessed with a laser confocal microscope (Carl Zeiss LSM780).

---

**Supplementary Figure 1.**
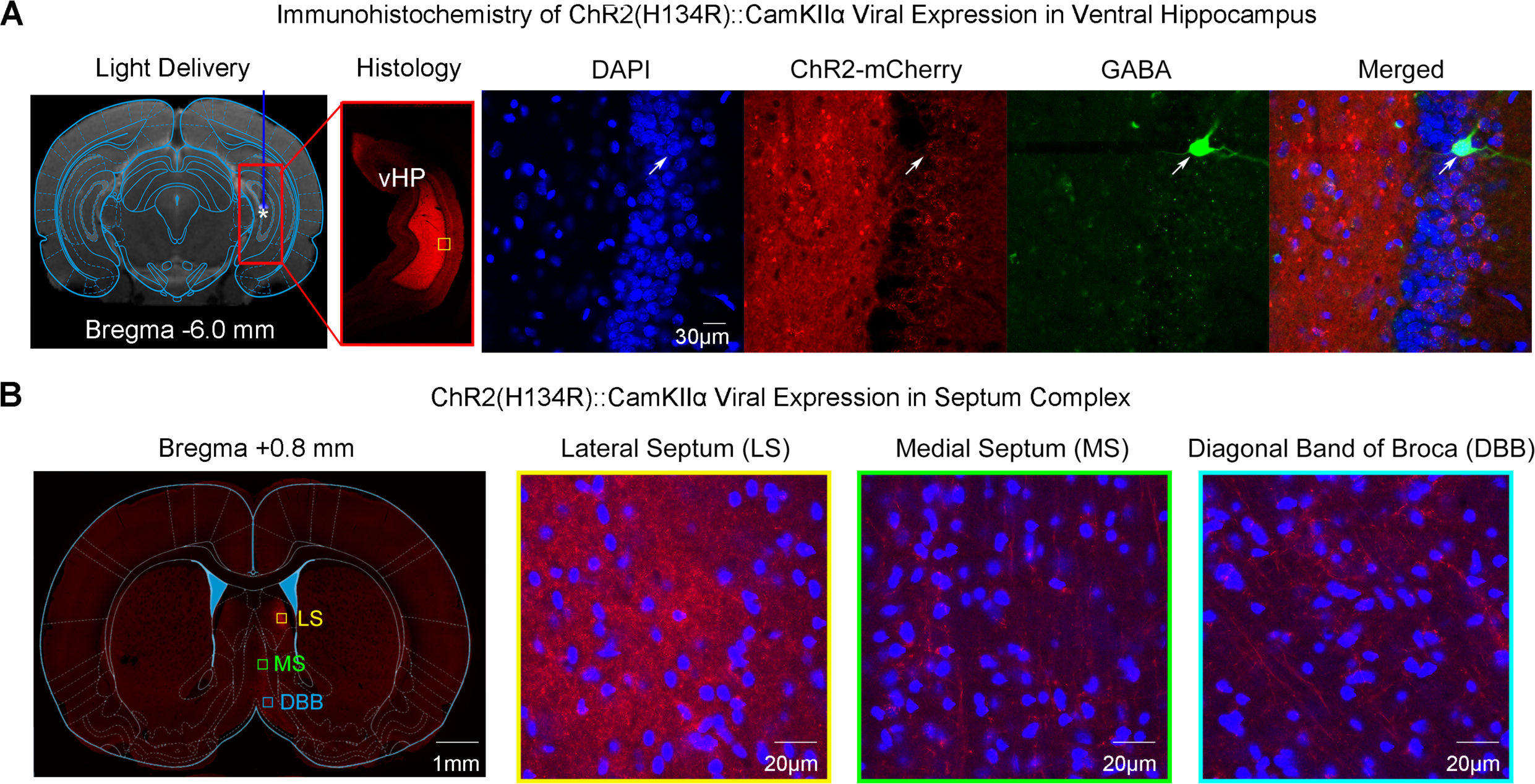
Histological characterization of ChR2::CaMKIIa viral expression in ventral hippocampus (vHP) excitatory neurons demonstrates no colocalization with vHP GABAergic neurons and their projection targets to the septum complex. ***(A)*** Confocal images of ChR2-mCherry expression in vHP; Lower magnification (*Left*) and higher magnification (*Right*). Overlay of images co-stained for the nuclear marker DAPI, mCherry, and inhibitory marker GABA revealed no colocalization between mCherry and vHP GABAergic neurons (indicated by white arrow). The yellow box indicates the location of magnified confocal images shown in *B*. ***(B)*** Confocal images of ChR2-mCherry expression in the septum complex. Lower magnification *(Left)* and higher magnification (*Right*). vHP projections synapse in the Lateral Septum (LS), Medial Septum (MS), and Diagonal Band of Broca (DBB), indicated by no colocalization between mCherry and DAPI.

**Supplementary Figure 2.**
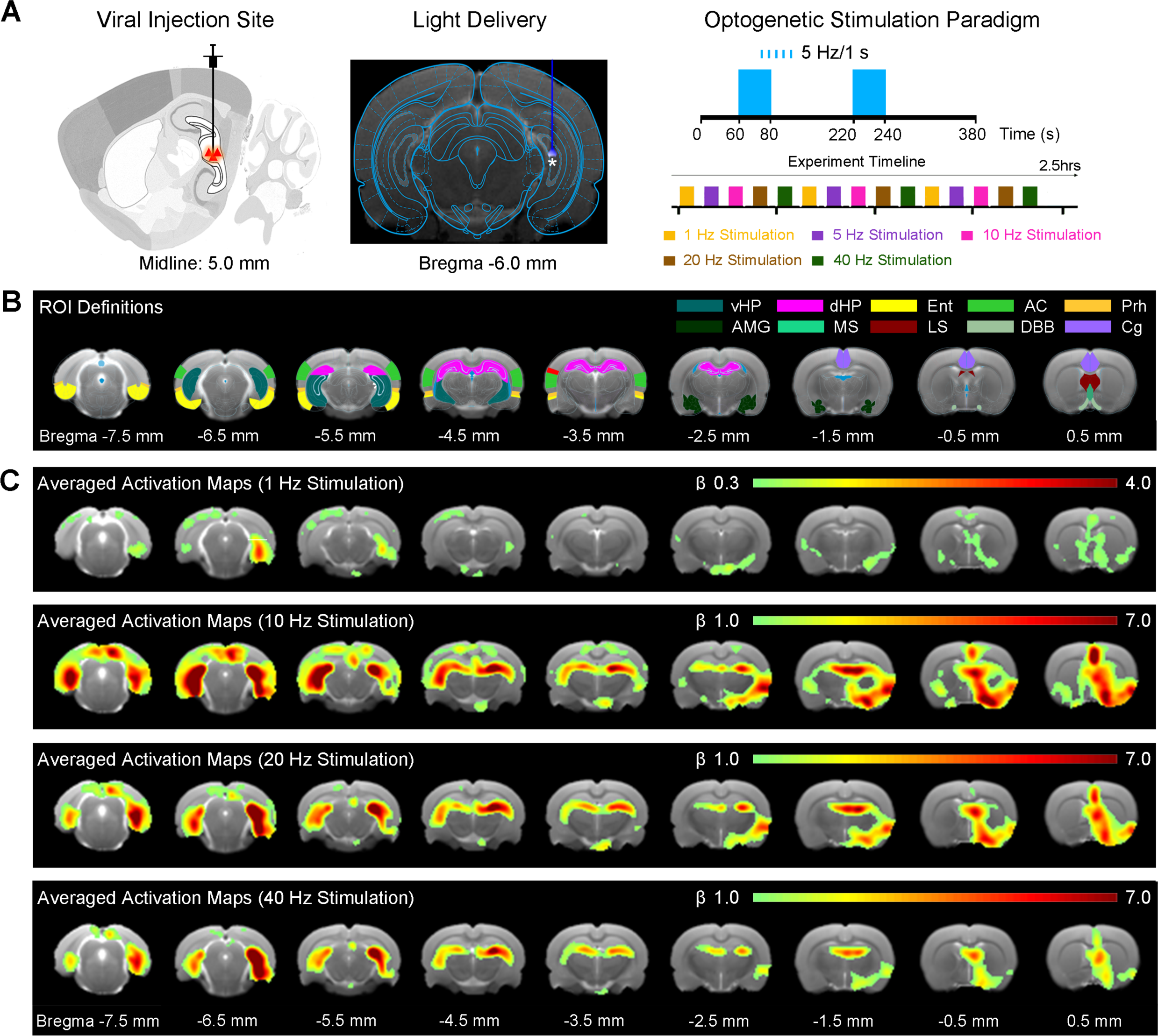
Brain-wide activations detected in the hippocampal formation, cortical and subcortical regions during optogenetic stimulation of excitatory neurons in vHP. *(A)* Schematic *(Left)* and T2-weighted anatomical image *(Middle)* shows the viral injection and fiber implantation sites, respectively. Optogenetic fMRI stimulation paradigm *(Right)*. Optogenetic stimulations were presented in a block-design paradigm (20 seconds ON; 140 seconds OFF). Five frequencies (1, 5, 10, 20, 40 Hz) were interleaved by trials. *(B)* Regions of interest (ROIs) defined by the rat brain atlas to extract the BOLD signal profiles. *(C)* Averaged activation (ȕ) maps of optogenetic stimulation of vHP excitatory neurons at 1, 10, 20, 40 Hz. Robust positive BOLD responses detected in bilateral vHP, dHP, Ent, AC, Prh, AMG, MS, LS, DBB, and Cg during 5 Hz optogenetic stimulation. Other frequencies (1 Hz, 10 Hz, 20 Hz, 40 Hz) evoked weaker BOLD responses in Prh, MS, LS, and Cg, while retained strong BOLD responses in vHP and dHP. (n = 6; t > 3.1, corresponding to p < 0.001). Abbreviations: Ventral Hippocampus (vHP); Dorsal Hippocampus (dHP); Entorhinal Cortex (Ent); Auditory Cortex (AC); Perirhinal Cortex (Prh); Amygdala (AMG); Medial Septum (MS); Lateral Septum (LS); Diagonal Band of Broca (DBB); Cingulate Cortex (Cg).

**Supplementary Figure 3.**
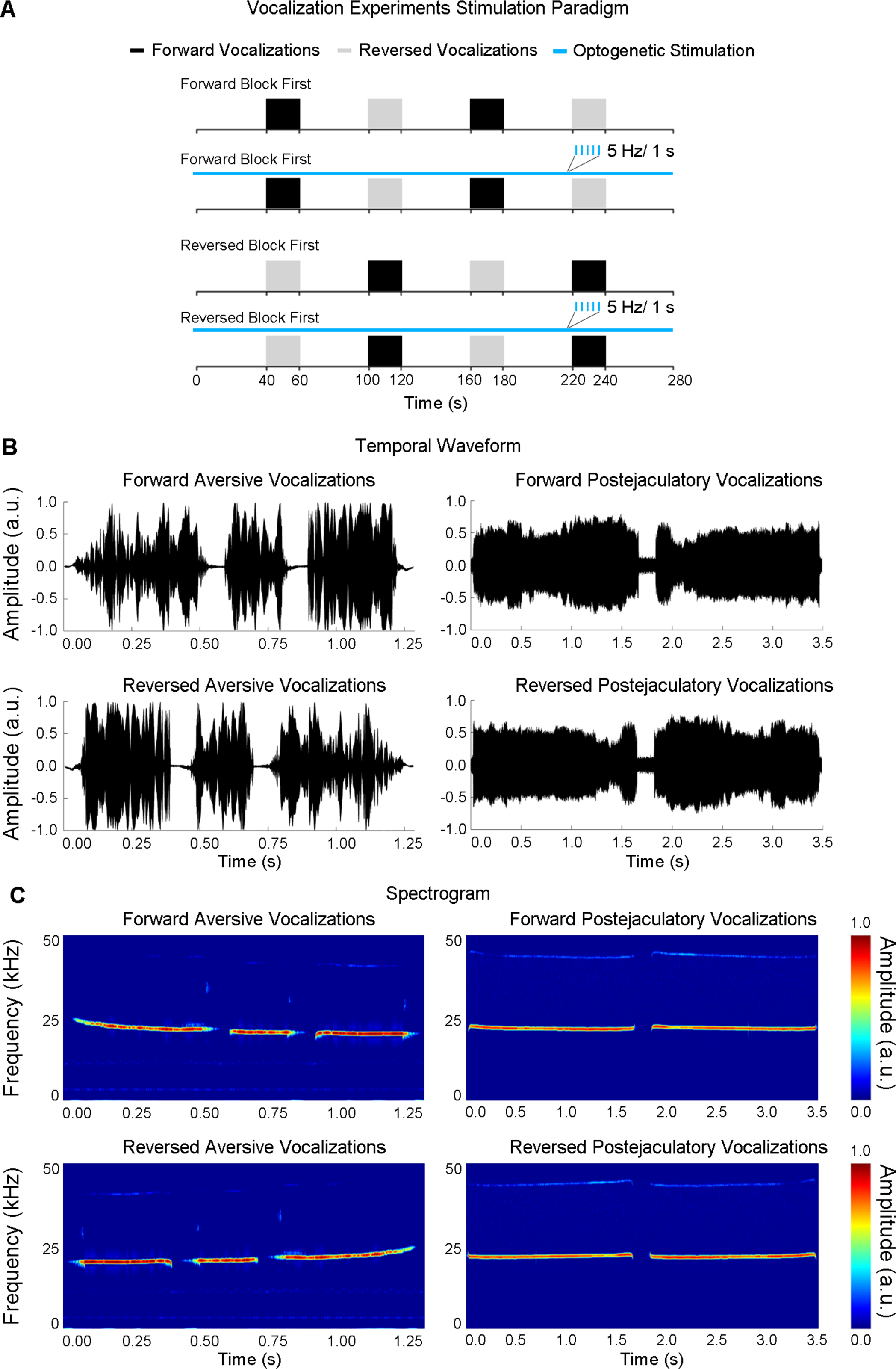
Auditory and optogenetic stimulation paradigms for the negative/aversive and positive/postejaculatory vocalizations experiments. *(A)* The standard block paradigm (20 seconds ON and 40 seconds OFF) was used to present rat vocalizations to the left ear. Forward and reversed vocalizations were interleaved during each auditory fMRI session. The paradigm was repeated six times for each animal, with the first block occupied by forward and reversed vocalization each three times, while continuous 5 Hz optogenetic stimulation was alternating between auditory fMRI sessions. *(B)* The temporal waveforms of forward *(top left)* and temporally reversed *(bottom left)* aversive vocalizations and forward *(top right)* and temporally reversed *(bottom right)* postejaculatory vocalizations. *(C)* The spectrograms of forward *(top left)* and temporally reversed *(bottom left)* aversive vocalizations and forward *(top right)* and temporally reversed *(bottom right)* postejaculatory vocalizations.

**Supplementary Figure 4.**
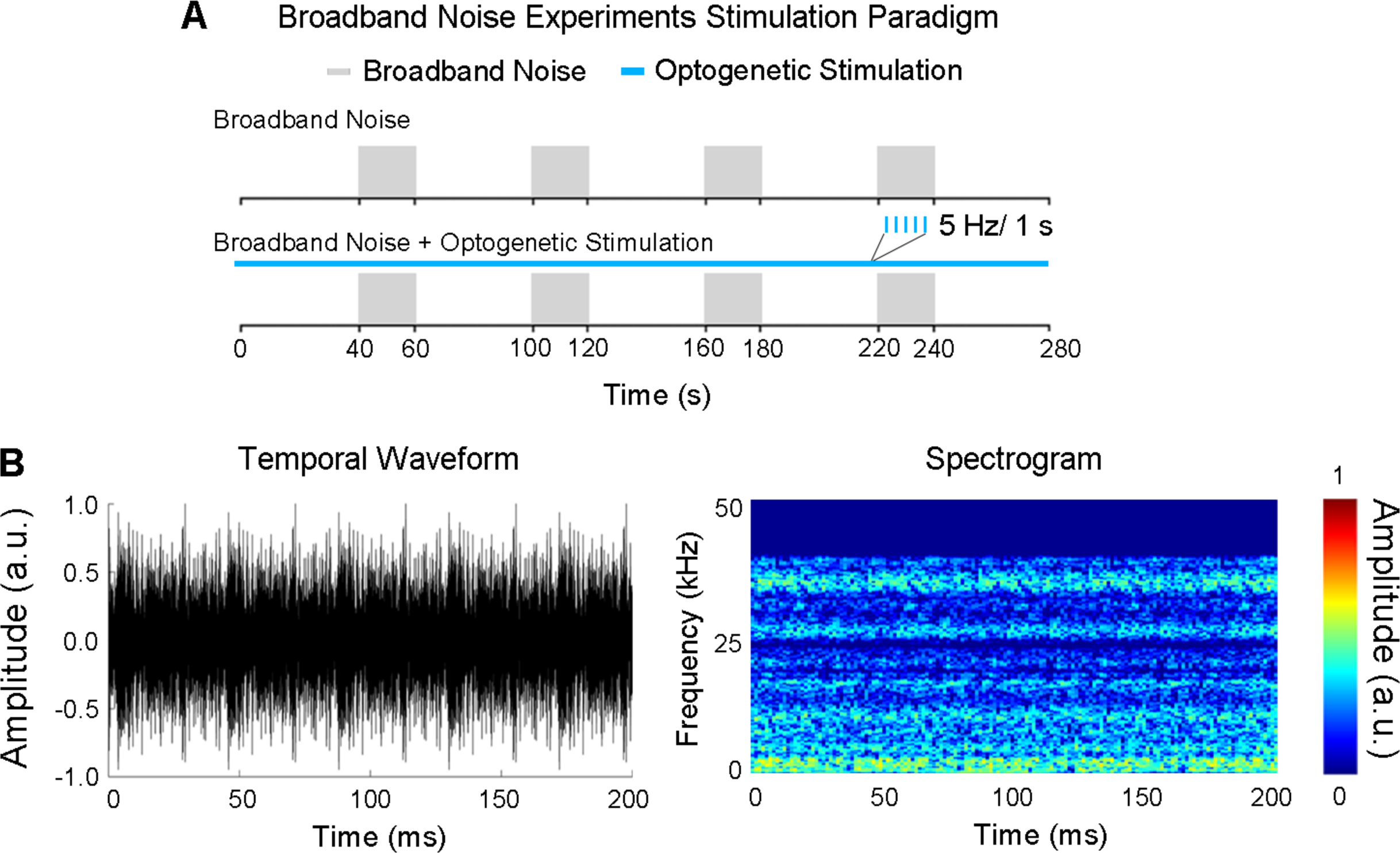
Auditory and optogenetic stimulation paradigms for broadband acoustic noise experiments. *(A)* The standard block paradigm (20 seconds ON and 40 seconds OFF) used to present broadband noise to the left ear, while continuous 5 Hz optogenetic stimulation was alternating between auditory fMRI sessions. *(B)* The temporal waveforms of the broadband noise. *(C)* The spectrograms of the broadband noise.

**Supplementary Figure 5.**
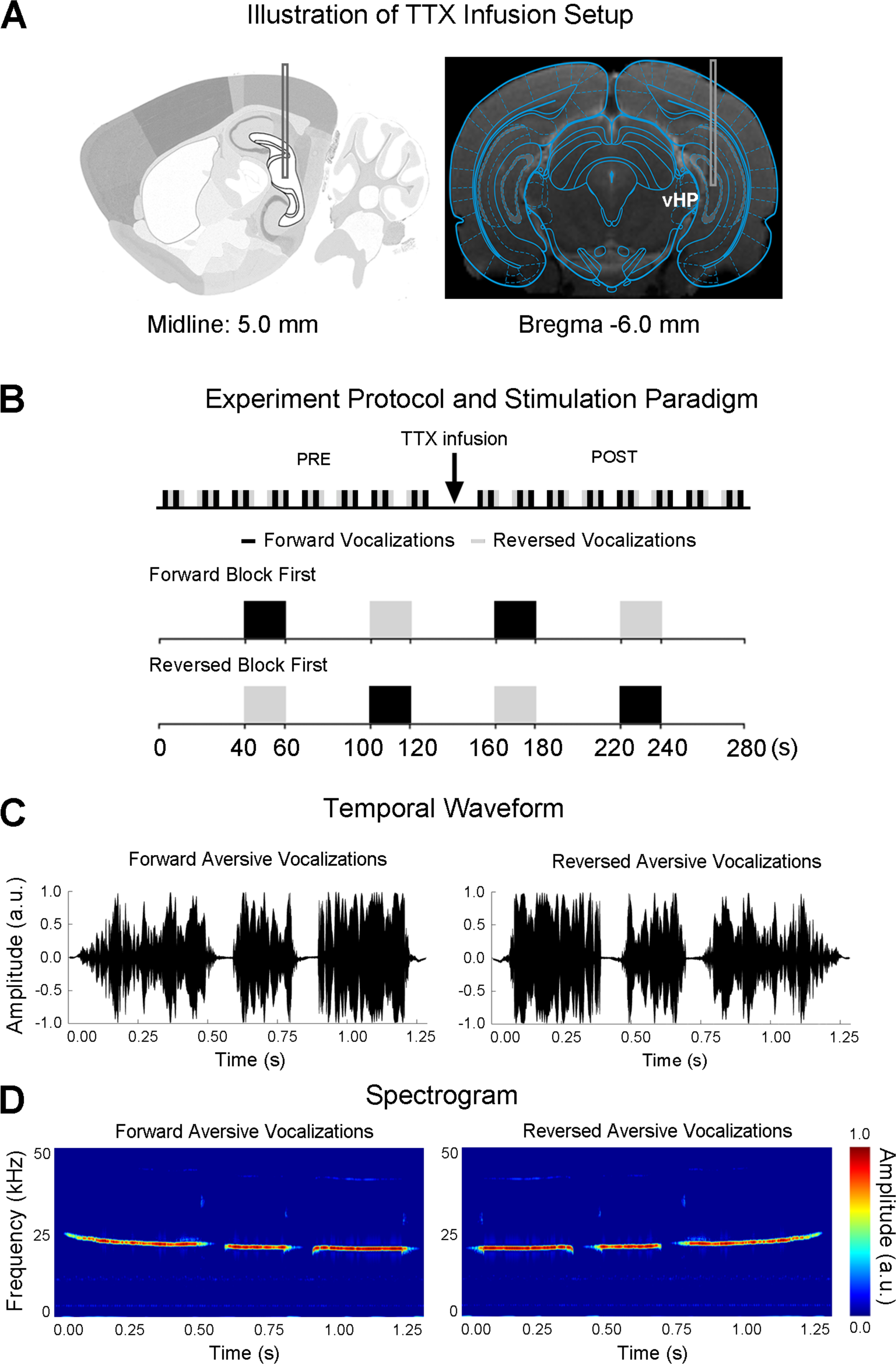
Illustration of tetrodotoxin (TTX) infusion to vHP, and corresponding auditory fMRI stimulation paradigm for the negative/aversive vocalization experiments. *(A)* TTX was infused to the vHP with an implanted cannula. *(B)* Sixteen fMRI sessions were typically acquired during an experiment. TTX was infused after eight fMRI sessions. Standard block design paradigm (20 seconds ON and 40 seconds OFF) was used to present vocalizations to the left ear. Forward and reversed vocalizations were interleaved during each auditory fMRI session. *(C)* The temporal waveform of forward *(left)* and temporally reversed *(right)* aversive vocalizations. *(D)* The spectrograms of forward *(left)* and temporally reversed *(right)* aversive vocalizations.

